# A T7 RNAP regulatory toolbox for cell-free network engineering and biosensing applications

**DOI:** 10.1101/2025.06.18.660355

**Authors:** Pao-Wan Lee, Seyed Saeed Mottaghi, Matthis Guillaume Lugnier, Sebastian J. Maerkl

**Affiliations:** Institute of Bioengineering, School of Engineering, École Polytechnique Fédérale de Lausanne, Lausanne 1015, Switzerland

**Keywords:** cell-free system, point-of-care diagnostic, gene regulation, transcription factor

## Abstract

T7 RNA polymerase is ubiquitously used in the fields of synthetic biology and biotechnology. Yet the ability to precisely and modularly regulate T7 RNAP remains surprisingly limited. Here, we developed a T7 RNAP regulatory toolbox consisting of programmable synthetic repressors, activators, and biosensors in a cell-free system. This toolbox enables the scalable design of T7 RNAP based gene regulatory networks and enables the rapid and sensitive detection of diverse biomolecules, including small-molecule drugs, antibodies, and proteins. By integrating a protein design pipeline, we generated biosensors using fully synthetic binders, demonstrating the potential for rapid development of novel protein-based sensors. We constructed a diagnostic cell-free system combining SARS-CoV-2 Spike protein sensing, gene regulatory based amplification, enzymatic amplification, and glucose based detection demonstrating the potential for point-of-care detection with high sensitivity. This work establishes a flexible and expandable framework for constructing gene circuits responsive to a wide range of biomolecules and demonstrates the potential for engineering point-of-care cell-free diagnostic assays.

## Introduction

Gene regulatory networks (GRNs) are central to biological information processing, enabling biological systems to sense, compute, and respond[1]. Synthetic biology has long pursued the bottom-up construction of GRNs to program cells and cell-free systems (CFSs) with user-defined functions, leading to hallmark circuits such as oscillators, toggle switches, synthetic pattern formation, and synthetic cell communication[2, 3, 4, 5, 6, 7].

Despite these advances, the scale and functionality of synthetic networks remain limited by their reliance on naturally derived regulatory parts. Many sensors and regulators are based on native transcription factors. As the demand for increasingly complex circuits grows, the limited availability of programmable, modular, and well characterized components has become a major bottleneck, hindering reliable and scalable GRN design[8].

In this work, we introduce a modular, and programmable toolbox for constructing synthetic transcriptional regulators including activators, repressors, and sensors with broad sensing capabilities. Our system is based on the ubiquitously used T7 RNA polymerase (T7 RNAP) and programmable zinc-finger (ZF) DNA-binding domains. All components are developed and characterized in the PURE CFS[9]. The open nature of CFSs enables rapid design–build–test cycles for gene regulatory elements, making it an ideal platform for prototyping and rapid deployment of individual components and complex circuits[10, 5]. The lack of cellular membranes in CFSs allows direct interaction between analytes and regulatory machinery, offering key advantages for diagnostic applications over cell-based approaches. CFSs are increasingly being applied in point-of-care (PoC) diagnostics due to their ease of use, low-cost, and compatibility with lyophilization, which enables transport without coldchain requirements and several CFSs have been developed for diverse targets, including pathogen RNA, metabolites, and heavy metals[11, 12, 13].

Our T7 RNAP regulatory toolbox achieved strong repression of T7 RNAP which is generally difficult to accomplish with native transcription factors. We also generated programmable T7 RNAP fusions which can be targeted to de novo promoters, drastically expanding the use of T7 RNAP in gene regulatory network construction with both activation and repression modules. We applied our toolbox to sense a wide range of biomolecular targets, from small-molecule drugs to proteins such as SARS-CoV-2 Spike and IgG antibodies in relevant clinical matrices. By integrating our platform with de novo protein design[14], we demonstrate a rapid workflow for generating synthetic biosensors using fully computationally designed binders. Finally, we constructed a diagnostic gene circuit that integrates Spike protein detection with gene circuit based amplification, enabling picomolar analyte detection using low-cost and readily available glucose meters for quantitative readout[15]. Together, these results highlight the potential of the synthetic T7 RNAP regulatory toolbox as a scalable, modular foundation for constructing novel gene circuits and sophisticated, high-performing, yet low-cost and easy-to-use biosensors.

## Results

### Strong and programmable T7 RNAP repression

T7 RNAP is widely employed in synthetic biology due to its high transcriptional activity, orthogonality to host machinery, and single-subunit architecture. Despite its utility, programmable repression of T7-driven transcription remains limited. Existing approaches rely heavily on naturally derived repressors such as LacI and TetR[16], which are insufficiently scalable for larger gene circuits. CRISPR-dCas9-based repression of T7-Lac promoters has been reported, but requires co-expression of LacI to maintain tight control over transcriptional activity[17]. Transcription activator-like effector (TALE) proteins have also been used to target the T7 promoter, with binding sites placed upstream or downstream of the core sequence. The observed repression was however modest (2–5 fold)[18]. To develop a repressor capable of strong and programmable inhibition of the T7 promoter, we established a streamlined workflow for rapid characterization of regulatory components using a CFS (Fig. 1a).

**Figure 1:**
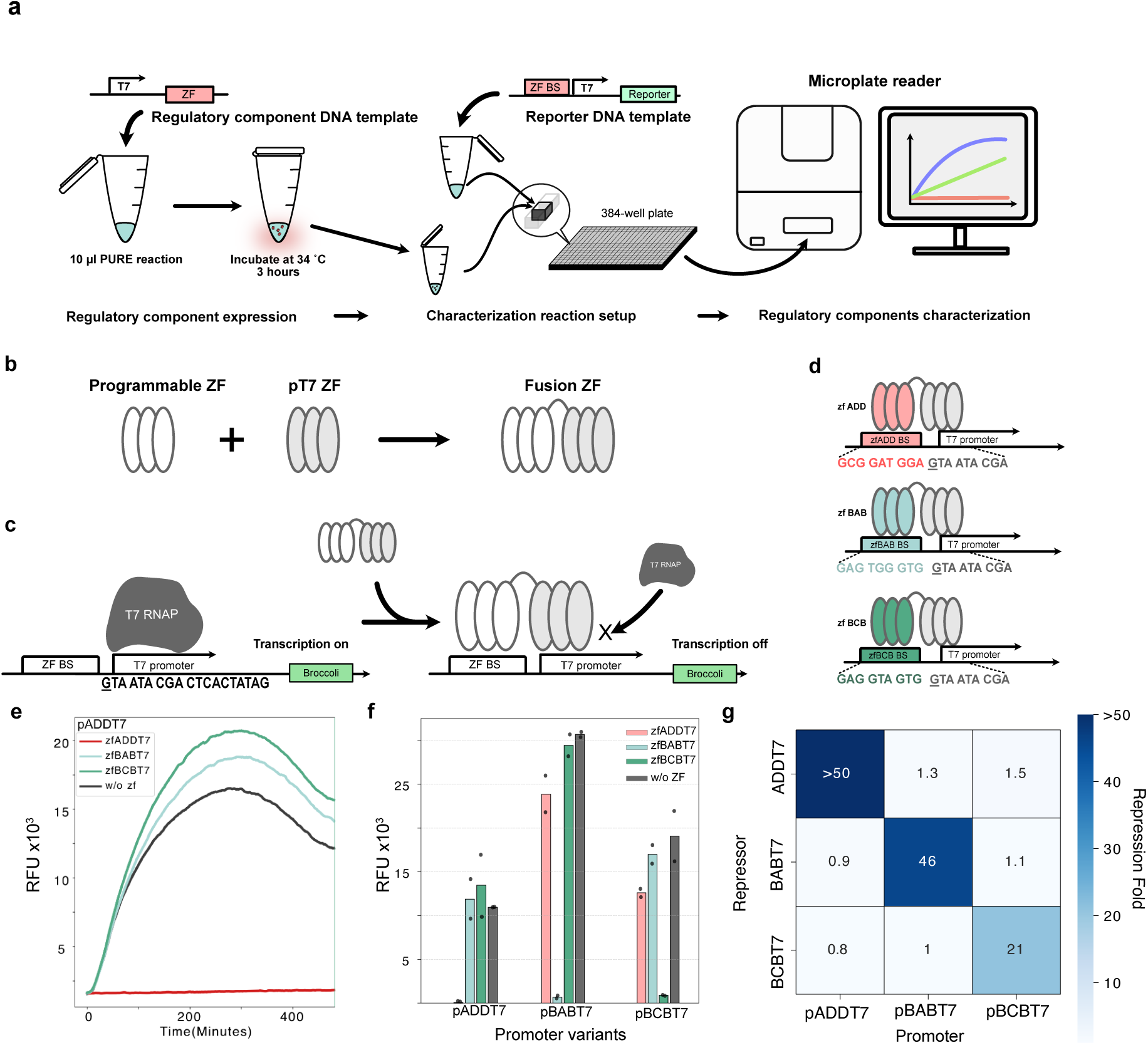
Characterization of regulatory components in CFSs: **a,** Workflow for characterizing gene regulatory components using the PURE system. Regulatory components are first expressed in a 10 µL PURE reaction at 34°C for 3 hours. Part of the reaction is then transferred into a second PURE reaction containing a reporter DNA template. Transcription kinetics is monitored in a 384-well plate using broccoli fluorescent RNA aptamer and a microplate reader. **b,** Design of the zfPR-zfT7 array for repressing T7 RNAP. **c,** Mechanism of repression by zfPR-zfT7 fusion proteins. The zfT7 domain targets a 9-bp DNA sequence overlapping the first 8 bp of the T7 promoter and an additional upstream G nucleotide, whereas the zfPR array targets a designed sequence upstream of the T7 promoter, providing increased repression strength and modularity. **d,** Three synthetic zfPR-zfT7 arrays and their corresponding promoters (pADDT7, pBABT7, and pBCBT7). **e,** Time-course of transcription from the pADDT7 promoter in the presence of different zfPR-zfT7 repressors showing specific and potent repression by the matched repressor (mean of n=2). **f,** Endpoint RFU (2 hours) of all promoter-repressor combinations, highlighting orthogonal repression behavior (mean of n=2). **g,** Repression fold calculated relative to the no-repressor condition with same promoter, showing strong and specific repression with minimal cross-reactivity (n=2).

Using this approach, we developed a programmable ZF-based repressor system that enables strong and orthogonal repression of the wild-type T7 promoter. This design builds on prior findings that tandem ZF arrays, when connected via appropriate linkers, can increase DNA binding affinity by up to 6,000-fold[19]. We engineered a fusion protein, zfPRzfT7 (Fig. 1b), consisting of a programmable ZF array targeting a user-defined 9 bp sequence (zfPR)[20], fused to a T7-specific ZF (zfT7) that binds to the GTAATACGA motif in the 5’ region of the T7 promoter (Fig. 1c). The zfT7 array was designed using established methods[21]. While zfT7 alone is unable to out-compete T7 RNAP for promoter binding, the addition of zfPR enabled the fusion protein to effectively compete with T7 RNAP and repress transcription.

To assess transcriptional repression, we used the RNA aptamer Broccoli in conjunction with the fluorophore DFHBI-1T as a fluorescent reporter for in vitro transcription[22]. We observed near-complete repression of the WT T7 promoter in the presence of the zfADDT7 repressor and its corresponding promoter (Fig. 1e). In contrast, reactions with non-cognate repressors exhibited transcriptional output comparable to the no-repressor control, confirming target specificity.

To further evaluate orthogonality and repression efficiency, we examined a 3×3 matrix of promoter–repressor pairs and calculated repression fold levels (Fig. 1f,g). The results demonstrate high orthogonality even among ZF consensus target sites differing by only three bases (e.g., zfBAB vs. zfBCB). Given that zfPR targets a 9 bp sequence, this system holds potential for the design of a large number of orthogonal T7 promoters and repressors. An additional advantage of this approach is that all promoter modifications are confined to the 5’ upstream region of the T7 promoter. As a result, the resulting transcripts retain identical 5’ UTRs, enabling predictable design by decoupling transcriptional regulation from translational variability.

### ZF guided T7 RNAP regulation

Although we demonstrated that our programmable repressors targeting the T7 promoter are highly functional, a programmable activator would also be highly beneficial for a comprehensive T7 RNAP gene regulatory toolbox. Since WT T7 RNAP exclusively recognizes the WT T7 promoter, it is not possible to achieve orthogonal gene activation without further modifications. Hussey et al. previously demonstrated that truncating the WT T7 promoter and inserting a ZF-binding site upstream (pZFd1T7 promoter) allows a ZF–T7 RNAP fusion protein to initiate transcription in a sequence-specific manner that is orthogonal to the native T7 promoter (Fig. 2a)[23]. By fusing distinct ZFs to T7 RNAP, transcription can be selectively activated from corresponding synthetic pZFd1T7 promoters. While this system was originally developed in *E. coli*, we successfully adapted it to CFSs, enabling rapid and modular characterization of activators and promoters under defined, tunable conditions.

**Figure 2:**
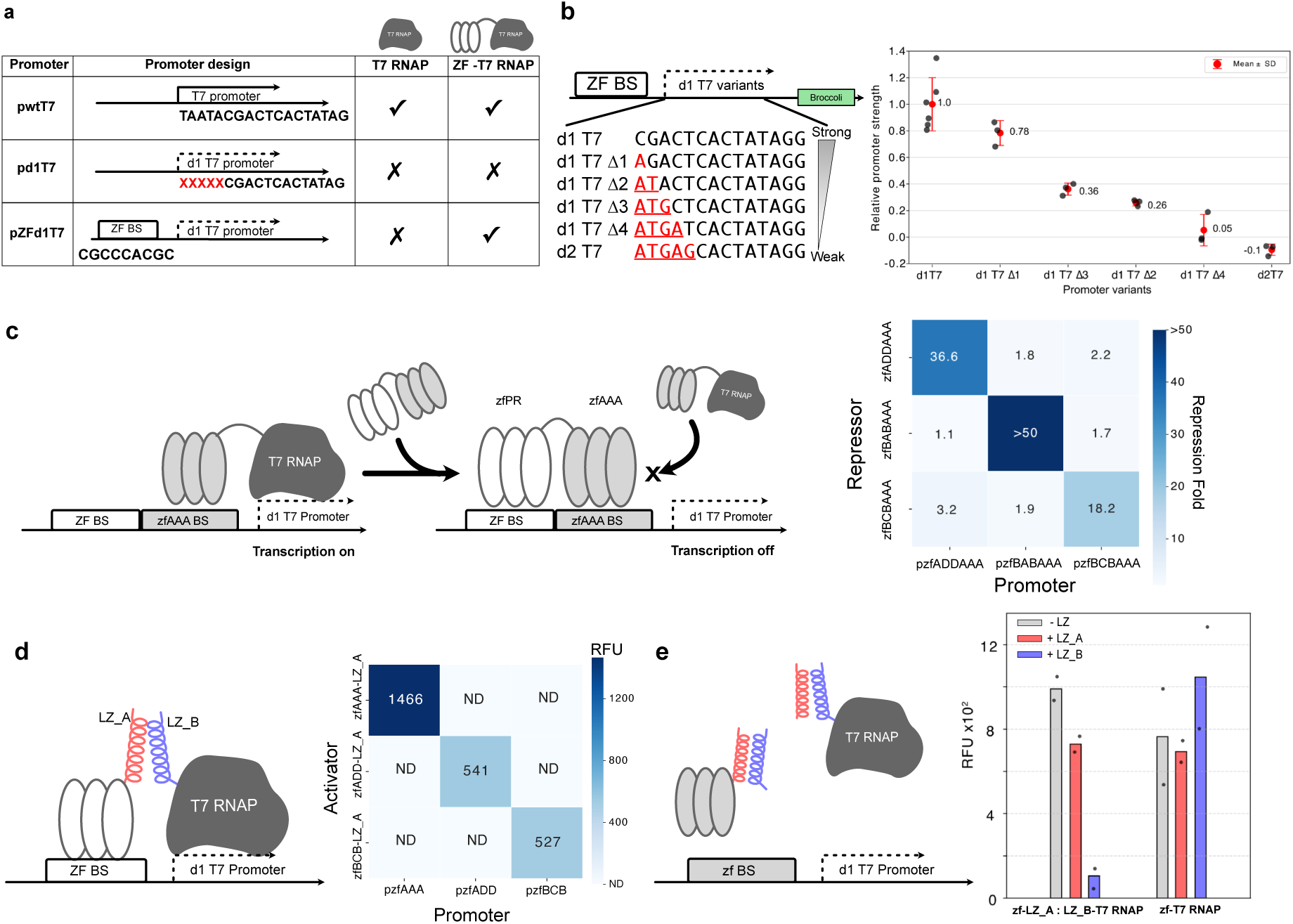
ZF guided T7 RNAP regulatory system: **a,** Table showing truncated T7 promoter, pd1T7, and pZFd1T7 with a modular ZF binding site upstream. **b,** pZFd1T7 promoter variants with different transcriptional strengths normalized by transcription activity of d1T7 promoter (d1T7 n=6, other promoter variants n=3, red dot indicates the mean, error bars indicate standard deviation). **c,** ZF-based repression of pZF(AAA)d1T7 promoter. zfAAA is fused to T7 RNAP, a modular zfPR array is fused to zfAAA as a repressor that strongly represses correpsonding pZFd1T7 promoters. Three repressor -promoter pairs were tested achieving repression levels of 18.2 to over 50 fold (n=2). **d,** ZF-based activation through LZ heterodimerization achieving activation with very low leakiness and off target effects (n=2). **e,** LZ-based repressors using LZ decoys where applied to interfere with the interaction between ZF and T7 RNAP (mean of n=2).

To enable tunable transcriptional output, we identified and characterized four variants of the pZF(AAA)d1T7 promoters with different strengths enabling modulation of expression levels, and improving the flexibility of gene circuit design (Fig. 2b). We also extended our repression strategy to the ZF–T7 RNAP activator platform. Initially, we observed that the zfAAA repressor alone was insufficient to repress transcription driven by the zfAAA–T7 RNAP fusion protein on the pZF(AAA)d1T7 promoter. We hypothesized that T7 RNAP retained residual affinity for the d1T7 promoter, stabilizing ZF–DNA binding and out-competing the repressor. To overcome this, we fused zfAAA to zfPR. The resulting zfPR-zfAAA repressor effectively repressed transcription from the pZF(AAA)d1T7 promoter, demonstrating strong repression with minimal leakiness (Fig. 2c).

Moreover, we expanded the activator system by employing ZFs from our characterized library[20]. Following the strategy described by Hussey et al.[23], we fused leucine zipper (LZ) heterodimer domains to both the ZF and T7 RNAP components, to activate specific transcription of pZFd1T7 promoters in a cell-free environment. This system provides a major advantage for multiplexed gene activation: a single LZ-T7 RNAP design can be recruited by different ZF-LZ activators through orthogonal LZ interactions. As shown in Fig. 2d, we observed no detectable crosstalk between different activators, indicating high orthogonality. Additionally, the LZ domains offer another layer of regulation. By introducing LZ decoys, we were able to inhibit activation, achieving approximately five-fold repression (Fig. 2e).

Together, this programmable approach for designing both activators and repressors with high orthogonality offers a promising foundation for constructing large-scale GRNs. The availability of well-characterized ZF libraries[20], as well as recent advances in deep learning–based ZF design[24], expands the potential of this toolbox for scalable and modular gene circuit engineering.

### T7 RNAP regulatory toolbox enables molecular sensing

To further expand and apply our T7 RNAP regulatory toolbox, we engineered ligand-induced dimerization mechanisms. Specifically, we developed various molecular sensing systems targeting a small molecule, antibody, or antigen. The presence of these molecules leads to ligand induced ZF-T7 RNAP transcriptional activation.

As a first demonstration, we developed a Venetoclax sensor based on a previously reported system where Venetoclax binding to BCL2 creates a new surface recognizable by designed protein binders[25]. By fusing BCL2 to a ZF and the DBVen1619 binder to T7 RNAP, we created a Venetoclax-inducible transcriptional activator (Fig. 3a). Venetoclax, a BCL2 inhibitor used to treat leukemia, has a narrow therapeutic window, and its plasma concentration should be closely monitored to balance efficacy and toxicity[26]. However, current clinical methods to monitor Venetoclax concentration rely on laborious liquid chromatography[27]. Our system offers a potentially faster and more cost-effective alternative for Venetoclax monitoring. We first assessed Venetoclax sensing in buffer solution achieving a limit of detection (LOD) of 30 nM (Fig. 3b), which was well within the range to detect under-dose levels. To assess clinically relevant performance of our system, we tested three Venetoclax concentrations corresponding to over-dose (10 *µ*M), therapeutic window (2 *µ*M), and under-dose thresholds (0.3 *µ*M) in human serum. 10% human serum was added to our cell-free reaction. The sensor was able to detect the 0.3 *µ*M under-dose threshold which corresponds to a 30 nM Venetoclax concentration in the final reaction volume, similarly to the result determined in simple buffer conditions. The sensor returned quantitative results clearly discriminating between the overdose, therapeutic window, and under-dose regimes (Fig. 3c) and the over-dose regime could be rapidly detected within 5 minutes (Figure S3a).

**Figure 3:**
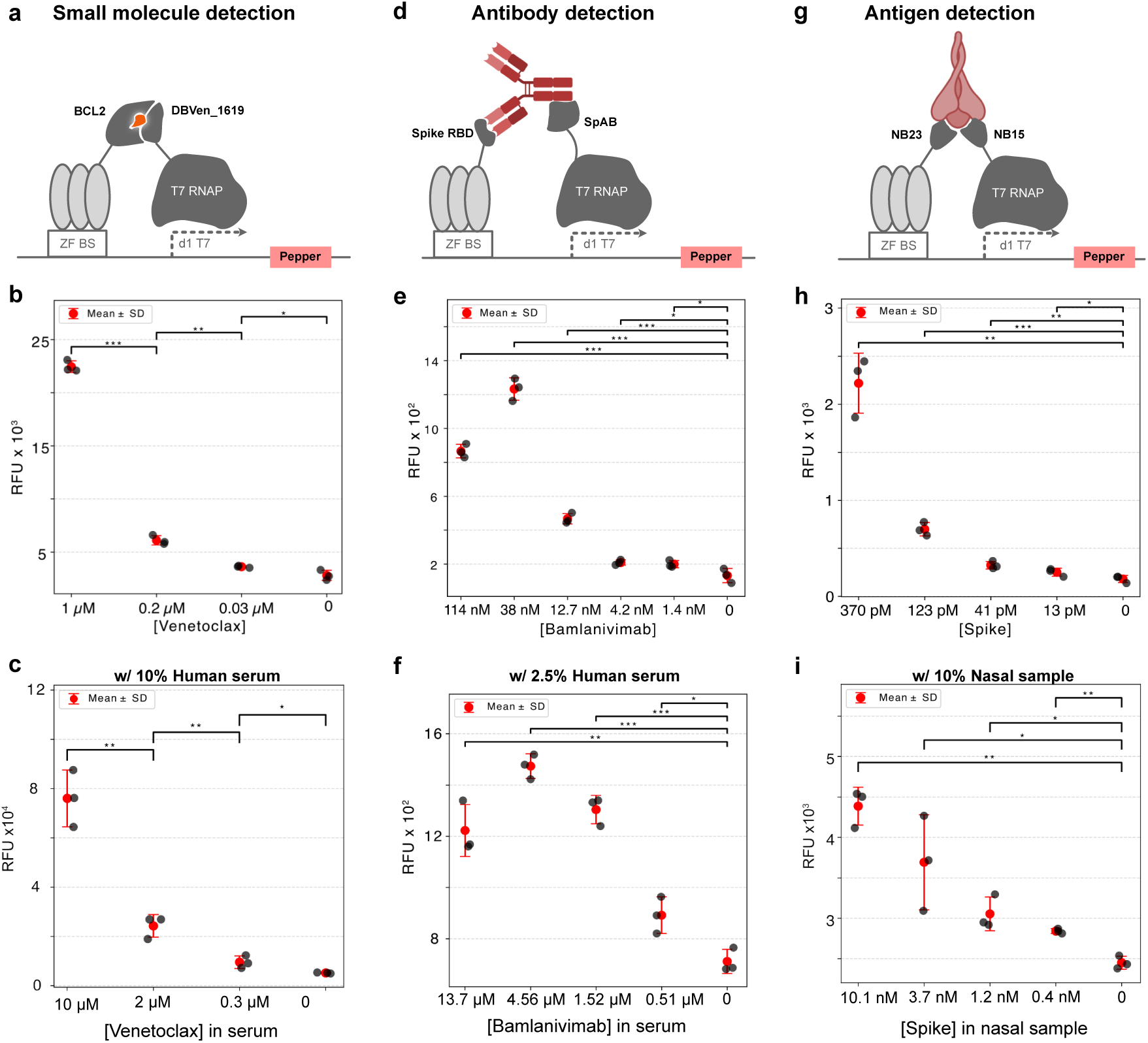
Modular biomolecules sensor of a small molecule and proteins: **a,** Venetoclax biosensor based on dimerization of de novo designed protein DBVen 1619 targeting the neosurface of the BCL2-venetoclax complex. **b,** Venetoclax detection levels in a standard PURE reaction. **c,** Detection of Ventoclax in human serum. Here Venetoclax concentrations indicate the concentration in the human serum matrix, not the final concentration in the cell-free reaction. **d,** Antibody biosensor detecting Anti-Spike IgG. This sensor relies on SpAB binding to the Fc region of IgG and Spike-RBD binding to the Fab region. **e,** Using Bamlanivimab as the analyte, the assay detects concentrations as low as 1.4 nM in standard PURE reactions. **f,** Detection of Bamlanivimab in human serum. Here Bamlanivimab concentrations indicate the Bamlanivimab concentration in the human serum matrix, not the final concentration in the cell-free reaction. **g,** SARS-CoV-2 Spike protein biosensor based on NB23 and NB15 nanobodies targeting two different epitopes on Spike. **h,** The system detects Spike concentrations as low as 13 pM in standard PURE reactions. **i,** Detection of Spike in nasal swab sample. Here Spike concentrations indicate the antigen concentration in the nasal sample matrix, not the final concentration in the cell-free reaction. (the following applies to all data panels: n = 3; red dot indicates the mean, error bars indicate standard deviation, * : p*<*0.05, ** : p*<*0.01, *** : p*<*0.001)

Next, we developed a human IgG antibody sensor based on antigen–antibody–SpAB interactions. In this system, the target antigen is recognized by the specific IgG antibody of interest, while the SpAB domain binds to the Fc region of the antibody. This interaction bridges the ZF and T7 RNAP fusion proteins, enabling antibody-dependent transcriptional activation (Fig. 3d). Using this design, we tested detection of Bamlanivimab, a therapeutic humanized IgG antibody used for treating SARS-CoV-2 infection. In standard cell-free reactions, we observed a concentration dependent response with an LOD of 1.4 nM (Fig. 3e). At high concentrations (114 nM) we observed a decrease in signal, which likely arises from the fact that at high concentrations individual antibodies are able to saturate both the antigen-ZF and the SpAB-T7RNAP fusion proteins.

In addition to antibody screening and characterization, the system could be employed in serological assays to monitor population infection rates during a pandemic, evaluate vaccine-induced immune responses, and track antibody titers over time. When tested in the presence of 2.5% human serum, the LOD increased to 12.7 nM, corresponding to ∼500 nM when back-calculated to the undiluted serum concentration (Fig. 3f). This detection limit is within clinically relevant ranges for detecting vaccine induced antibody titers and potentially infection induced titers as well. The use of the SpAB domain which recognizes the Fc region of IgGs is currently limiting the use of the sensor in serological applications, as the SpAB domain binds to all IgG antibodies present in a serum sample. To improve the sensor for these type of applications one could substitute the SpAB domain with another antigen domain which would result in detection of only antigen specific antibodies.

Lastly, to assess the possibility to use our sensing concept for clinically relevant antigen detection we constructed a Spike protein sensor using two nanobodies (NBs) binding distinct epitopes on the SARS-CoV-2 spike protein[28]. By fusing NB23 to ZF and NB15 to T7 RNAP, we achieved transcriptional activation in the presence of Spike (Fig. 3g). When testing Spike protein directly in the cell-free IVT reaction we were able to achieve very low detection levels of 13 pM (Fig. 3h). When testing the system’s sensitivity using 10% nasal swab samples the sensor achieved an LOD of 40 pM, corresponding to an effective Spike concentration of 400 pM in the original sample (Fig. 3i).

In summary, our platform demonstrates broad versatility: by simply exchanging the recognition modules, we successfully detected a diverse range of biomolecules, including small molecules, antibodies, and viral proteins in complex matrices such as human serum and nasal swab samples. These results highlight the system’s potential for low-cost, modular, and field-deployable diagnostic applications.

### Computational protein design coupled with CFSs enables rapid sensor development

Traditional biosensing methods often rely on antibodies or nanobodies, which require laborintensive screening and optimization, thereby slowing the development of new diagnostic tools. Recent advances in de novo protein design including RFdiffusion, MaSIF-seed, and BindCraft, have opened new avenues for creating protein binders tailored to specific molecular targets[14, 29, 30]. We sought to leverage de novo binder design coupled with cell-free systems to demonstrate the possibility of rapidly developing a molecular sensing system. In this study, we aimed to streamline sensor development by integrating a computational binder design pipeline into our platform, replacing the NBs previously used in our Spike protein detection system with fully de novo designed binders.

The binding regions of NB23 and NB15 were assigned as hotspots in the publicly available RFdiffusion pipeline to design two de novo binders targeting Spike RBD (Fig. 4a). The design process was performed on a single RTX 4080 GPU. In two weeks we generated over 100’000 protein designs (2430 structures with 32-64 sequence variants per structure) of which we selected the 45 top-scoring designs for testing. No experimental high-throughput screening or affinity maturation of these designs was performed. Top-ranking designs were tested for function using a rapid cell-free screening assay eliminating any time-consuming cloning steps. Each binder was fused to a ZF domain and co-expressed with a reporter gene in a 10 *µ*L cell-free reaction containing 10 nM Spike protein. The top-performing binder for each binding epitope was then validated in triplicate reactions.

**Figure 4:**
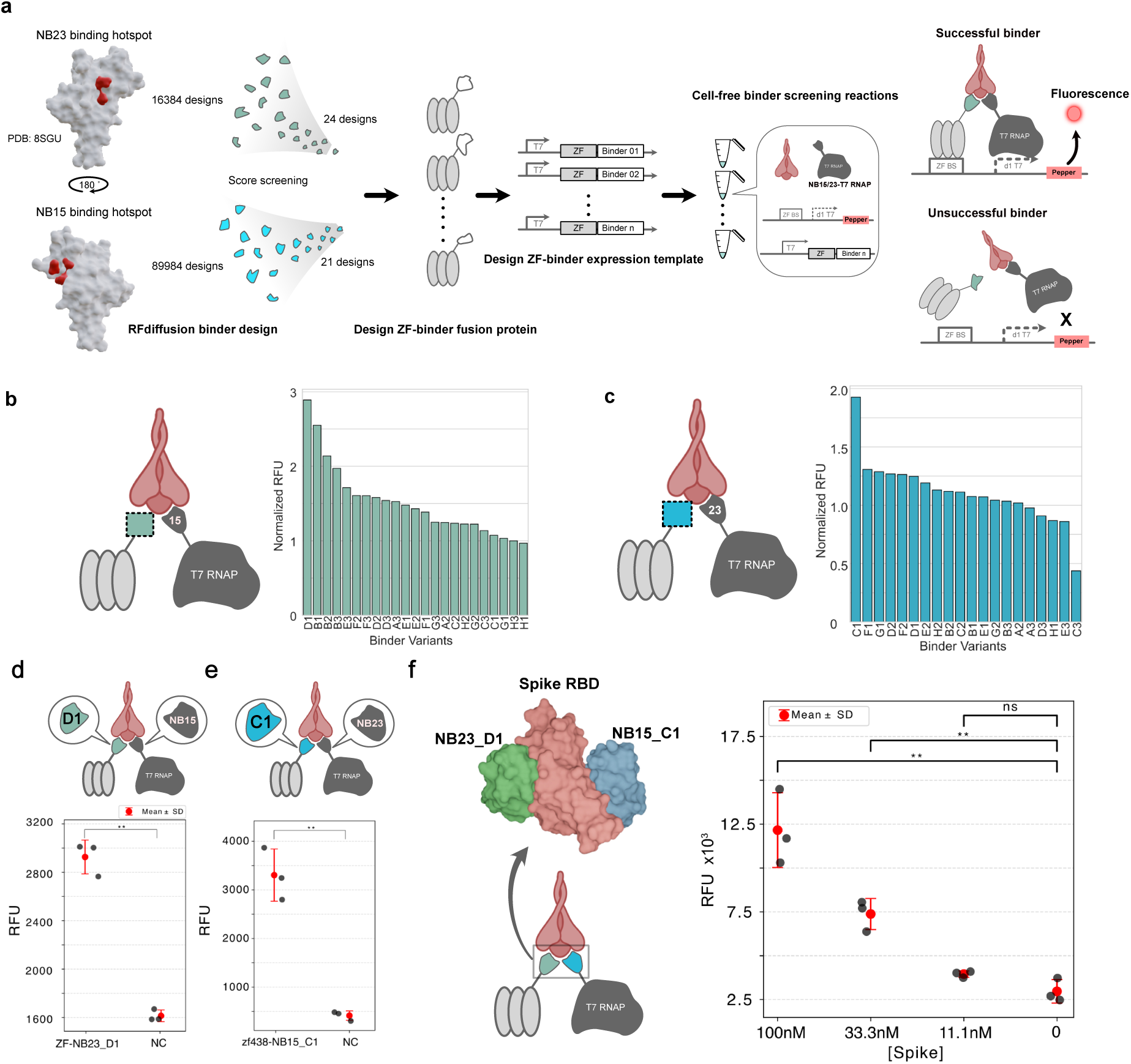
Rapid de novo binder design and screening: **a,** Workflow for de novo binder design and screening. Binding epitopes of NB23 and NB13 on Spike RBD were used to define binding hotspot residues as input for RFdiffusion to generate candidate binders. Top-scoring designs were fused to ZF and synthesized as linear DNA templates for rapid in vitro screening. Cell-free reactions included the ZF-binder expression DNA template, NB-T7 RNAP fusion proteins, Spike RBD, and a fluorescent reporter. Successful binders reconstitute ZF-Spike-T7 RNAP trimerization, activating transcription and producing fluorescence. **b,** Screening results for binders targeting the NB23 hotspot residues. Fluorescence intensities are normalized to the negative control (n=1). **c,** Screening results of binders targeting the NB15 hotspot. Fluorescence intensities are normalized to the negative control (n=1). **d,** Validation of the top-performing ZF-NB23 D1 variant (n = 3; red dot indicates the mean, error bars indicate standard deviation; formatting applies to panels e and f). **e,** Validation of the top-performing ZF-NB15 C1 variant (n=3). **f,** Construction of a fully de novo designed Spike protein sensor using NB23 D1 and NB15 C1 binders (n=3). The de novo sensor detects Spike antigen down to 33.3 nM (n=3). (* : p*<*0.05, ** : p*<*0.01, *** : p*<*0.001)

We successfully identified two de novo binders capable of functionally replacing NB23 and NB15 respectively, which we refer to as NB23 D1 and NB15 C1 (Fig. 4b-e). Remarkably, this was achieved with a small in silico design campaign followed by experimentally testing a small number of designs: 24 for NB23 and 21 for NB15. We attribute this high success rate in part due to recent improvements in de novo protein design methodologies as well as to the specific characteristics and attributes of our sensing platform. Unlike classical molecular detection methods such as ELISA, which require high-affinity and stable complexes between binder and analyte in order to withstand wash steps and achieve good performance, our system appears to tolerate binders with low or moderate dissociation constants (Kd) which is likely the case for the two de novo binders developed here. This is because in our system no stable complexes need to be generated, and even low affinity, transient interactions between binder and analyte are likely sufficient to give rise to complex assembly and consequent transcription in our system.

After testing the two de novo designed binders individually we constructed a fully de novo designed Spike sensing system using both discovered binders. Although the use of the two de novo binders resulted in a marked decrease in LOD, they were nonetheless capable of detecting Spike concentrations as low as 33.3 nM (Fig. 4f), which outperforms a recent study reporting an LoD of 150 nM using classical nanobodies in combination with a split T7 RNAP system[31].

These results demonstrate that de novo protein binders can be rapidly generated and seamlessly integrated into our system. Moreover, the system enables rapid and cost-effective binder screening without the need for labor-intensive cloning, display, and sequencing methods. We performed de novo binder design without the use of any specialized GPU computing cluster platform, although the use of high-throughput computing clusters would drastically reduce the in silico binder design process from 1-3 weeks to 1-3 days. Continued improvements in computational binder design will likely enable high-affinity binders to be developed with increased ease, allowing highly sensitivity molecular diagnostic assays to be rapidly developed in conjunction with our cell-free screening and detection approach.

### Cell-free point-of-care molecular diagnostics

CFSs are appealing platforms for developing next-generation PoC molecular diagnostic assays [11, 12, 13]. Particularly the open nature of CFSs (lacking physical boundaries such as cell-membranes), their potential for integrating sophisticated molecular sensing, the possibility to produce these system at low-cost, and the ability to lyophilize the reactions for extended shelf-life and distribution at ambient temperature, makes them promising platforms for the development of PoC assays. To create a PoC compatible system, we integrated a low-cost and simple quantitative readout method based on ubiquitously available glucose readers[32]. We integrated trehalase as the downstream reporter, which converts trehalose into glucose, into our Spike protein sensing system creating a simple-to-use one-pot reaction (Fig. 5a). The nasal sample containing spike protein is added directly to the cell-free reaction, and glucose levels are measured after a one hour incubation period (Fig. 5b). We optimized the NB15-T7 RNAP concentration to reduce background activity and determined the optimal incubation time for glucose readout(Fig. S5). Using this approach, we were able to detect Spike protein in nasal sample at concentrations as low as 3.33 nM (Fig. 5c).

**Figure 5:**
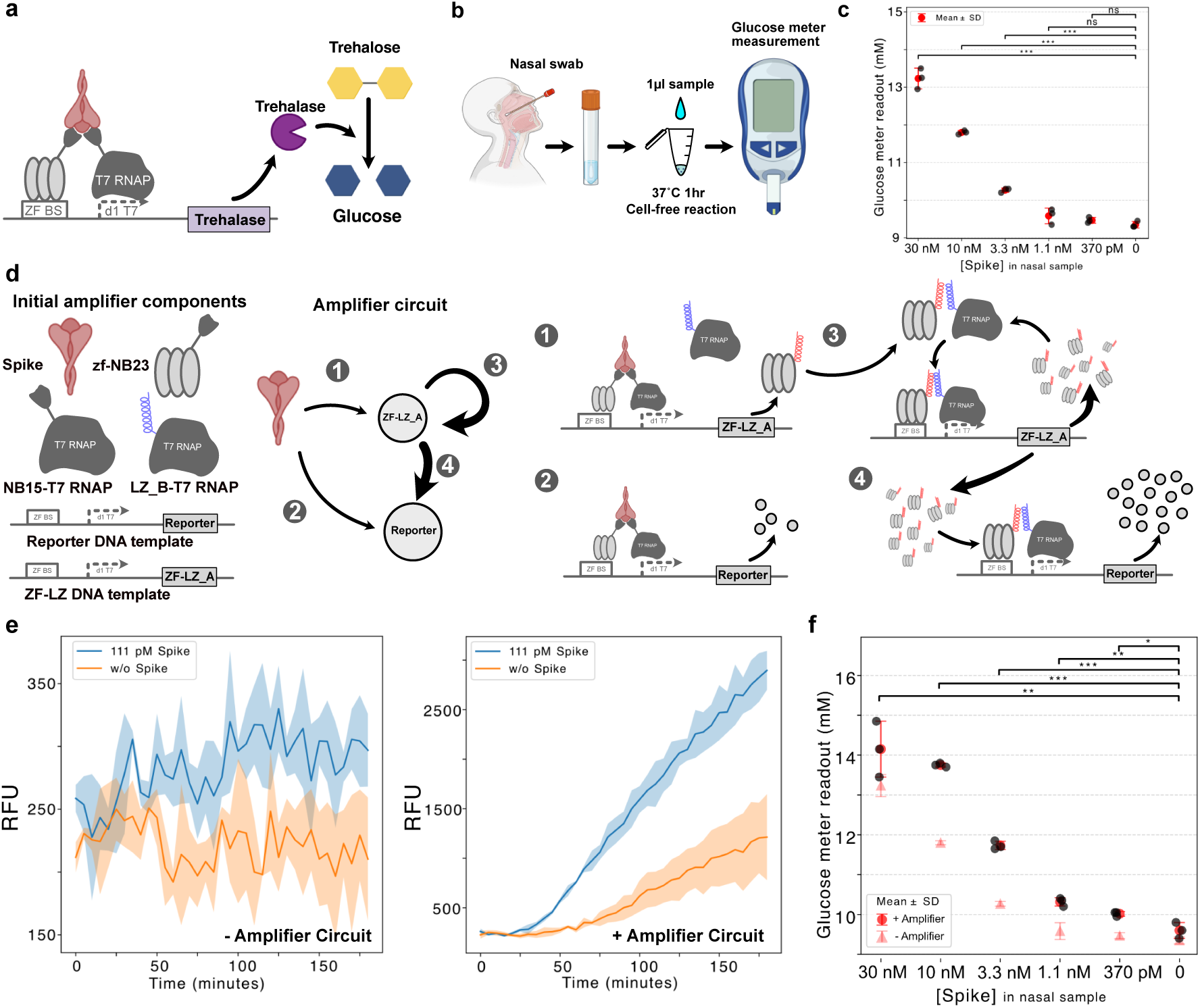
Point-of-care diagnostic system development: **a,** Schematic of the spike protein sensing system coupled to trehalase expression. Spike protein induces trehalase production, which converts trehalose into glucose for detection via a commercial glucose meter. **b,** Workflow of the point-of-care diagnostic. 1 µL of sample containing varying concentrations of spike protein is added to 9 µL of the cell-free reaction. After 1 hour of incubation at 37°C, glucose levels are quantified using a glucose meter. **c,** Glucose endpoint measurements after 1 hour of incubation, showing achieving an LOD of 330 pM. (n = 3; red dot indicates the mean, error bars indicate standard deviation; formatting applies to panels f) **d,** Design of the amplification circuit. The system uses a self-activating transcriptional activator ZF-LZ A. In the presence of spike antigen, ZF-LZ A is expressed, which binds to LZ B-T7 RNAP and further drives its own expression, amplifying the response under low spike concentrations. **e,** Reactions with and without the amplifier circuit using Pepper fluorescent reporter. The reaction with amplifier circuit shows significantly enhanced and distinct transcriptional activity. (n = 3, the curves represent the mean, and shaded regions indicate standard deviation). **f.** Integration of the amplifier circuit with glucose readout. The addition of the amplifier lowers the LOD by an order of magnitude compared to the system without amplifier.

While the trehalase - glucose reader combination provided a convenient PoC compatible detection method, its LOD was higher than that achieved using the Pepper RNA aptamer reporter. But CFSs have the unique capacity to perform sophisticated signal amplification without increasing assay complexity. To enhance sensitivity, we designed an amplifier gene circuit using our regulatory components (Fig. 5d). The circuit comprises two additional elements: (i) a DNA template encoding ZF–LZ A under the control of a pZFd1T7 promoter, and (ii) a LZ B–T7 RNAP fusion protein. Spike protein is sensed as described previously by activating transcription of the trehalase reporter gene, but it now also can activate transcription of ZF-LZ A, which dimerizes with LZ T7 RNAP giving rise to a positive feedback loop. As the reporter gene is driven by the same promoter, this loop amplifies reporter expression.

To evaluate the amplifier circuit, we monitored transcription kinetics using the Pepper fluorescent RNA reporter. At 111 pM spike protein, the original system lacking the amplifier yielded low fluorescence near background levels, whereas the amplifier circuit generated significantly stronger and easily distinguishable responses (Fig. 5e). We next integrated this amplification system into the trehalase - glucose meter assay. With 10% nasal swab samples, we achieved an LOD of 37 pM spike protein in the reaction, corresponding to 370 pM in the original sample improving the LOD by almost an order of magnitude compared to the system lacking the amplifier module (Fig. 5f).

These results highlight the modularity and scalability of our platform. By combining modular sensor with enzyme-mediated signal conversion and an amplifier gene circuit, we demonstrate how complex systems can be rapidly designed and assembled from our T7 RNAP regulatory toolbox for sensitive PoC diagnostic.

## Discussion

Regulatory components form the basis of gene circuit design. T7 RNAP in particular is one of the most commonly used RNA polymerases in synthetic biology, cell-free synthetic biology, and biotechnology, primarily due to it’s strong transcriptional activity. One caveat of T7 RNAP’s strong transcriptional activity is that it has proven difficult to repress, nor do regulatory elements akin to sigma factors exist capable of recruiting T7 RNAP to different promoters. We developed a comprehensive T7 RNAP regulatory toolbox which relies on ZF DNA binding domains which are small protein-domains that can be readily expressed in CFSs. By designing a double array of ZF domains, with one array directly competing for the T7 RNAP promoter, we were able to achieve high fold repression. Modularity is achieved by modifying the second ZF array which enables targeted repression of specific promoters. To extend the system we demonstrated that an ablated T7 RNAP promoter is also functional and versatile in CFSs[23], allowing the creation of sigma factor like activation functions, which can be repressed either in cis through transcription factor - DNA binding, or in trans by disrupting dimerization. We thus established a comprehensive cell-free T7 RNAP toolbox using a set of small, well-behaved protein domains that can be readily expressed and lead to minimal resource loading. The use of ZFs allows the creation of a vast number of possible DNA binding specificities by using either existing ZF libraries[20] or AI driven design[24].

In addition to serving as a versatile platform for cell-free synthetic circuit engineering, our approach is well-suited for the development of advanced biosensing and molecular diagnostics applications. We demonstrated that the system can be rapidly engineered to detect not only proteins but also small molecules. We successfully quantitated the small-molecule drug Venetoclax at clinically relevant concentrations in human serum. The platform also enables the detection of protein targets, such as antibodies and antigens (SARS-CoV-2 spike protein), supporting applications in antibody characterization, and protein engineering. By integrating computational protein design with cell-free screening, we showed that novel biosensing or molecular diagnostic assays can be rapidly developed.

PoC diagnostic tests are crucial for decentralized, low-cost testing to monitor disease or environmental monitoring. The most widely used format has been lateral flow assays, which offer simplicity but are limited in flexibility and sensitivity. In contrast, CFSs are emerging as powerful new alternatives for PoC applications [11, 12, 13]. To demonstrate that our system is compatible with PoC applications we integrated the expression of a sugar converting enzyme, which allows quantitative readouts to be obtained with simple, low-cost, and widely available glucose meters. We also showed that more sophisticated functionalities such as gene regulatory based circuit amplifiers can be integrated into the system. We were able to detect 370 pM of spike protein in nasal sample using this approach (corresponding to 37 pM of Spike in the reaction mixture) and detection was possible in 1 hour or less. For comparison, a recently published report using a split T7 RNAP system was able to detect 750 nM Spike protein (corresponding to 150 nM in the reaction mixture) using a qualitative filter paper approach [31]. Our approach is thus 3-4 orders of magnitude more sensitive and returns quantitative results.

This T7 RNAP regulatory toolbox and CFSs engineering approach spans a wide range of potential application and use cases ranging from gene regulatory circuit engineering, molecular sensing, protein screening, and molecular diagnostics. In the basic research domain, the toolbox enables rapid gene regulatory circuit engineering using robust and well characterized parts. It also provides the exciting possibility to directly integrate sensing functionalities, which could give rise to sophisticated feedback control systems capable of sensing small molecules and proteins. The rapid progress in computational protein design methods will further enhance the ability to create novel functions and applications. We demonstrated the possibility of integrating the toolbox with downstream gene regulatory networks and it is also possible to integrate complex RNA based circuits in cases were it is advantageous to retain a transcription-only system. Even multiplexed molecular assays should be within reach, as well as the possibility to perform molecular computation in a one-pot cell-free PoC reaction. This gives rise to exciting possibilities in engineer high-performing molecular PoC diagnostic tests.

## Methods

### Plasmids and linear DNA template

Plasmids for the expression of PURE system proteins were kindly provided by Yoshihiro Shimizu (RIKEN Center for Biosystems Dynamics Research, Japan). Plasmids encoding binder–T7 RNAP variants were cloned into the pQE30 vector and transformed into NEBExpress® Iq Competent *E. coli* (NEB, C3037I) for protein expression. The plasmid encoding ZF-NB23 was cloned into pET-21(+) and transformed into *E. coli* BL21(DE3) for expression. Linear DNA templates were synthesized by Twist Bioscience. The design of these templates is detailed in Supplementary Table S1. Gene fragments used for cell-free reactions were separated by agarose gel electrophoresis, and the desired bands were excised and amplified via PCR using Q5^®^ High-Fidelity 2X Master Mix (NEB, Cat. No. M0492) and adapter primers. The PCR products were purified using a commercial kit (Zymo Research, Cat. No. D4014). The sequences of the gene fragments are listed in Supplementary Table S3.

### Protein expression and purification

For PURE proteins expression, the same process was used for all proteins [33]. For expression and purification of binder-T7 RNAP variants, after the plasmids are transformed to NEBExpress® Iq Competent *E. coli*, single colonies from the transformation were picked and inoculated into a small volume (5-10 mL) of LB medium containing 100 *µ*g/mL ampicillin (Amp) as a preculture. The preculture was incubated at 37°C overnight at 260 RPM. Overnight cultures were then used to inoculate larger volumes of LB medium (500-1000 mL) containing 100 *µ*g/mL Amp at a 1:100 ratio. The cultures were incubated at 37°C and 260 RPM for two hours, after which protein expression was induced by adding isopropyl *β*-D-1-thiogalactopyranoside (IPTG) to a final concentration of 0.1 mM, followed by an additional three-hour incubation at 37°C and 260 RPM. Cells were harvested by centrifugation at 4°C and 3,220 × g for 10 min and stored at −80°C. For protein purification, buffers were prepared according to Supplementary TableS4), and tris(2-carboxyethyl)phosphine (TCEP) was added just before use. The pellets were resuspended in 15 mL of buffer A and lysed by sonication using a 130-watt probe with the following parameters: 4 × 20 s pulse on, 20 s pulse off, and 70% amplitude. The lysate was then centrifuged at 21,130 × g for 20 min at 4°C to remove cell debris. Since all the proteins were expressed in the soluble phase, the supernatant was loaded onto ∼3 mL of IMAC Sepharose 6 Fast Flow beads (GE Healthcare, catalog no. GE17-0921-07). Washing and elution steps were performed using buffers containing 25 mM (23.75 mL of buffer A + 1.25 mL of buffer B) and 450 mM imidazole (0.5 mL of buffer A + 4.5 mL of buffer B), respectively. The eluted protein was then buffer-exchanged either by overnight dialysis with dialysis tubing cellulose membrane (Sigma-Aldrich, catalog no. D9652) against 1 L of HT buffer followed by 3 hours of dialysis against Stock buffer A under shaking at 150 RPM at 4°C, or by using a 15 mL Amicon Ultra Filter Unit −3 kDa MWCO (Millipore, catalog no. UFC9003) with centrifugation at 3,220 × g and 4°C. Before this, the sample was diluted with HT buffer 6 times, followed by the addition of the corresponding volume of Stock buffer B. The protein concentration was measured by Nanodrop based on absorbance at 280 nm. Some precipitation might be observed after dialysis with HT buffer, but after dialysis with Stock buffer A, the precipitation would decrease. The remaining precipitation can be removed with syringe filter. For expression and purification of ZF-binder, we used the same procedure, except including 20 *µ*M of ZnSO_4_ in LB media, purification and stock buffers.

### Time-course florescent measurement

Cell-free reactions are loaded into 384-well black microplate with transparent bottom (Corning,catalog no. 3544) for fluorescent measurement. The measurements were conducted with a plate reader (BioTek Synergy H1 Multimode Reader). For Broccoli aptamer with DFHBI-1T: excitation 485 nm, emission 515 nm. For Pepper-HBC620: excitation 577 nm, emission 620 nm. The temperature was set to 34 °C for regulatory components characterization reactions.

### Zinc finger repressor and promoter design

The programmable ZF were selected from previous characterized ZF library[20]. The zinc finger that targets pwtT7 was designed with the standardized ZF design method[21]. We used a default ZF backbone amino acid sequence, and used ZF that specifically target GTA, ATA and CGA from the published library. The linker sequence between zfPR and zfT7 is from the publication, which has shown to increase the binding affinity significantly [19].

ZF proteins bind DNA in an antiparallel orientation: the N-terminal finger of the protein recognizes the 3’ end of the target DNA sequence, while the C-terminal finger recognizes the 5’ end. To repress the WT T7 promoter, the ZF binding site was placed immediately upstream of the T7 promoter. The zfPR was fused to the C-terminus of the zfT7 to achieve repression.

For ZF-T7 RNAP, the ZF was fused to the N-terminus of T7 RNAP. Given the antiparallel binding orientation—where the N-terminal finger binds to the 3’ end and the C-terminal finger to the 5’ end—the target DNA sequence for the ZF was designed as the reverse complement of the intended recognition sequence. This ensures proper ZF binding orientation and positions T7 RNAP correctly toward the downstream d1 T7 promoter. The same principle applies in designing the ZF-guided T7 system repressors. The target sequence for the ZF repressor is placed upstream of the zfAAA site and is the reverse complement of its recognition sequence. To construct the ZF repressor in ZF-T7 RNAP system, zfPR domain was fused to the N-terminus of zfAAA, enabling transcriptional repression.

### Characterization of regulatory components

For characterization of the WT T7 repressor, repressors were first pre-expressed using PUR-Express (NEB, catalog no. E6800) and then diluted 1000-fold prior to characterization. A 0.5 µL of the diluted protein was added to 8.5 µL of reaction mix, yielding a total volume of 9 µL. Reactions were transferred to a 384-well plate and incubated at room temperature for 10 minutes. Transcription was initiated by adding 1 µL of reporter DNA. Reactions were then incubated at 34 °C, and Broccoli fluorescence was measured every 2 minutes using a plate reader. All reactions contained 10 µM DFHBI-1T for activation of the Broccoli RNA aptamer. RFUs were averaged from two timepoints (±2 minutes) centered at 2 and 122 minutes. Each condition was performed in duplicate, and the average of replicates was used to calculate repression fold, defined as the ratio of RFUs from reactions lacking ZF to those including ZF.

For characterization zfxxxAAA repressors, a similar procedure was followed. Pre-expressed repressors were diluted 250-fold, and 0.5 µL was added to 8.5 µL of reaction mix. After a 10-minute pre-incubation at room temperature, 1 µL of reporter DNA was added. Broccoli fluorescence was monitored as described above. Due to a decrease in signal during the initial 18 minutes, RFUs at 18 minutes were used as the starting point and 138 minutes as the endpoint. Repression fold was calculated as described above.

To characterize activation from the pZFd1T7 promoter using zfPR–LZ A activators, pre-expressed activators were diluted 100-fold. The measurement time interval was set to 5 minutes. RFUs were averaged from two timepoints (±5 minutes) centered at 5 and 125 minutes. For LZ-based repressors, it followed the same procedure as the activators, but with pre-expressed repressors diluted 20-fold before adding to reactions. The analysis window was also from 5 to 125 minutes, using a ±5-minute averaging window.

To characterize pZF(AAA)d1T7 promoter variants with different strengths, IVT was used instead of PURExpress, as the only required protein was the zfAAA–T7 RNAP fusion, eliminating the need for additional expression steps. Promoter strength was quantified by normalizing the RFU yield of each variant to that of the pZF(AAA)d1T7 promoter. RFU yield was calculated as the difference in fluorescence between 2 and 62 minutes (±2 minutes), averaged across replicates. Each variant was tested in triplicate, except pZF(AAA)d1T7, which was measured in six replicates.

### Nasal sample collection

Nasal samples were collected following standard procedures for commercial rapid tests. Both nostrils were swabbed using the same swab, making ten circular motions along the nasal walls. The swab was then stirred ten times in 500 *µ*L of PBS.

### Biomolecule sensing reactions

Biomolecule sensing reactions were conducted using the PURExpress supplemented with pre-expressed ZF–binder and purified binder–T7 RNAP fusion proteins. The Pepper RNA aptamer was used as the fluorescence reporter in combination with 10 µM HBC620 fluorophore, which provides reduced background signal compared to the Broccoli/DFHBI-1T system. All sensing conditions were tested in triplicate across varying analyte concentrations.

The yield of RFU was calculated as the difference between endpoint and starting point fluorescence signals, with each value averaged over two adjacent timepoints to reduce noise. The specific timepoints used for each experiment are detailed below. For statistical analysis and LOD determination, Welch’s unpaired one-sided t-test was used to compare RFU yields between analyte-added and negative control conditions. Significance thresholds were defined as follows: (* : p*<*0.05, ** : p*<*0.01, *** : p*<*0.001)

For Venetoclax sensing, the ZF–BCL2 fusion protein was pre-expressed in PURExpress at 34°C for 3 hours and added to 10 µL sensing reactions containing the remaining components. In conditions containing 10% human serum, Murine RNase Inhibitor (NEB, catalog no. M0314) was added at 1 U/µL to prevent RNA degradation. RFU yield was calculated as the difference between values at 2 and 62 minutes (averaged ±2 minutes).

For antibody sensing, ZF–RBD was pre-expressed in PURExpress at 34°C for 3 hours with the addition of PURExpress® Disulfide Bond Enhancer (NEB, catalog no. E6820S) to assist in proper folding of the RBD domain. The pre-expressed ZF–RBD was then added to 10 µL reactions to detect bamlanivimab. In samples containing 2.5% human serum, Murine RNase Inhibitor was again included at 1 U/µL to preserve RNA integrity. RFU yield in serum-containing conditions was calculated as the difference between 2 and 62 minutes. In serum-free conditions, an initial decline in fluorescence was observed during the first 20 minutes; thus, the analysis window was adjusted to 22–82 minutes.

For spike protein sensing, reactions were carried out using an in vitro transcription (IVT) format. Reactions contained purified ZF–NB23, NB15–T7 RNAP, reporter DNA template, and 5× IVT buffer (see Supplementary Table S5). Defined concentrations of spike protein were added to 20 µL IVT reactions to determine the system’s LOD. In samples containing 10% nasal sample, Murine RNase Inhibitor was again included. RFU yield was calculated as the difference between signals at 2 and 62 minutes (±2 minutes).

### De novo spike protein binder design and screening

PDB 8sgu file is applied as input to RFdiffussion using the following Google colab notebook. Epitopes that recognized by NB23 and NB15 are first identified and provided as hotspot for RFdiffusion pipeline design. For the NB23 replacement binder, amino acid residues 340, 354, 457 were selected as hotspot. 7 rounds of design were conducted, 16384 binders are generated and 24 binders are selected for testing. For the NB15 replacement binder, amino acid residues 381,428,517 were selected as hotspots, 16 rounds of design were conducted, 89984 binder were generated and 21 binders were selected for in vitro screening. The RFdiffusion pipeline was conducted with a single GeForce RTX 4080 GPU. The general design parameters were as follows: contig size was 75-100, iterations between 50-200 (100 iterations were used predominantly), 32-256 designs with 64 or 32 sequence variants per design, symmetry = none, partial T = auto, use beta model = true, mpnn sampling temp = 0.1 or 0.2, rm aa = C, use solubleMPNN = false, initial guess = true, num recycles = 3, use multimer = false.

DNA templates encoding ZF–binder fusion proteins were prepared at a stock concentration of 10 ng/µL. For screening, a working solution was prepared at 0.1 ng/µL by diluting the stock. Binder screening reactions were assembled using the master mix detailed in Table S6. To screen binders intended to replace NB23, the master mix included NB15–T7 RNAP; conversely, to screen binders intended to replace NB15, NB23–T7 RNAP was included in the mix. Each reaction had a total volume of 10 µL, including 1 µL of DNA template encoding a different ZF–binder fusion variant.

Fluorescence resulting from expression of the Pepper reporter was monitored at 2-minute intervals using a microplate reader. Fluorescence intensities were normalized to a negative control lacking ZF–binder DNA. The binder variant showing the highest normalized fluorescence was selected for validation. Validation experiments were performed in triplicate, and statistical significance was assessed using a t-test as described in the previous section.

Once both binders were validated, they were fused to ZF and T7 RNAP, respectively, to construct a fully synthetic binder pair for Spike protein detection. The LOD was determined by calculating the difference between the fluorescence signal at 62 minutes (averaged over ±2 minutes) and the background signal at 2 minutes (also averaged over ±2 minutes). Statistical significance was assessed using the t-test procedure described previously.

### Homemade PURE preparation

Due to the leakiness caused by high concentration of T7 RNAP in commercial PURE systems, we prepared our homemade ΔT7 RNAP PURE, which contains the other 35 PURE proteins except T7 RNAP. By using homemade ΔT7 RNAP PURE and optimizing NB15T7RNAP and ZF438-NB23 concentrations, we minimized leakiness. The homemade PURE and homemade ΔT7RNAP PURE protein mixtures were prepared by combining the individual purified proteins. For the homemade ΔT7 RNAP PURE protein mixture, T7 RNAP was excluded. The concentration of all 36 proteins are prepared based on previous published paper [34].

### Glucose meter readout reaction

The reaction contained Solution A from PURExpress as the energy solution, along with homemade ΔT7RNAP PURE proteins and commercial ribosome (NEB, catalog no. P0763). These components were added to 10 *µ*L reactions at final volumes of 4 *µ*L, 1.3 *µ*L, and 0.45 *µ*L, respectively, or at the same relative proportions in reactions of other volumes. Trehalose (Sigma-Aldrich, catalog no. T9531) was included at a final concentration of 10 mM. The pZFd1T7-tre37A template, NB15-T7RNAP, and ZF438-NB23 were used at final concentrations of 20 nM, 10 nM, and 50 nM, respectively. For reactions with the amplifier system, the pZFd1T7-ZF438-LZ template and T7RNAP-LZ were added at final concentrations of 3 nM and 10 nM, respectively (Supplementary Table Table S7). For the evaluation of the amplifier circuit with the Pepper aptamer, we prepared a similar reaction, replacing trehalose with HBC620 at a final concentration of 10 nM, and using reduced concentrations of the templates pZFd1T7-ZF-LZ A and pZFd1T7-Pepper, at 1 nM and 6 nM, respectively (Supplementary Table S8). For glucose measurement, the reaction mixtures were incubated at 37°C in a thermocycler, and 1 *µ*L samples were taken at different time intervals. Glucose concentration was measured using a glucose meter (Ascensia Diabetes Care, Contour XT) by directly loading the sample onto the strip. Different spike concentrations were prepared by diluting the recombinant trimeric SARS-CoV-2 spike protein (LubioScience, catalog no. LU2010) into the nasal sample or PBS, and 1 *µ*L of each was used in 10 *µ*L reactions, or in the same relative proportion for reactions of other volumes. In negative controls, the same volume of nasal sample or PBS was added to the reaction instead of the spike protein. For spike titration reactions in triplicate, after 1 hour of incubation at 37°C, the samples were stored on ice to stop the reaction. The glucose concentration of each triplicate was measured at least twice, and the average of the measurements for each triplicate was used for statistical analysis and plots. In the reactions with nasal sample, 1 unit/*µ*L of Murine RNase Inhibitor was also included.

## Author Contribution

P.W.L., S.S.M. and M.G.L performed experiments. P.W.L. and S.S.M., and S.J.M. designed experiments, analyzed data, and wrote the manuscript.

## Acknowledgements

This work was supported by a Swiss National Science Foundation Sinergia Grant (514558), and a Swiss National Science Foundation MINT grant (514337). Part of the graphical schematic representations are created using BioRender.com. We thank Ali Mekki Berrada, Elisa Sarah Cabos and Maxence Christian Pierre Calamand for their valuable discussions on the amplification circuit.

## Conflict of interest

The authors declare that a patent application related to the biosensor technology described in this manuscript has been filed.

## References

[1] Alon, U. An Introduction to Systems Biology: Design Principles of Biological Circuits (Chapman and Hall/CRC, 2006).

[2] Elowitz, M. B. & Leibler, S. A synthetic oscillatory network of transcriptional regulators. Nature 403, 335–338 (2000).

[3] Gardner, T. S., Cantor, C. R. & Collins, J. J. Construction of a genetic toggle switch in escherichia coli. Nature 403, 339–342 (2000).

[4] Basu, S., Gerchman, Y., Collins, C., Arnold, F. & Weiss, R. A synthetic multicellular system for programmed pattern formation. Nature 434, 1130–1134 (2005).

[5] Niederholtmeyer, H. et al. Rapid cell-free forward engineering of novel genetic ring oscillators. eLife 4 (2015).

[6] Swank, Z., Laohakunakorn, N. & Maerkl, S. J. Cell-free gene-regulatory network engineering with synthetic transcription factors. Proceedings of the National Academy of Sciences 116, 5892–5901 (2019).

[7] Adamala, K. P., Martin-Alarcon, D. A., Guthrie-Honea, K. R. & Boyden, E. S. Engineering genetic circuit interactions within and between synthetic minimal cells. Nature Chemistry 9, 431–439 (2016).

[8] Brophy, J. A. & Voigt, C. A. Principles of genetic circuit design. Nature Methods 11, 508–520 (2014).

[9] Shimizu, Y. et al. Cell-free translation reconstituted with purified components. Nature Biotechnology 19, 751–755 (2001).

[10] Lee, P.-W. & Maerkl, S. J. Regulatory components for bacterial cell-free systems engineering. ACS Synthetic Biology 13, 3827–3841 (2024).

[11] Pardee, K. et al. Rapid, low-cost detection of zika virus using programmable biomolecular components. Cell 165, 1255–1266 (2016).

[12] Voyvodic, P. L. et al. Plug-and-play metabolic transducers expand the chemical detection space of cell-free biosensors. Nature Communications 10 (2019).

[13] Gräwe, A., et al. A paper-based, cell-free biosensor system for the detection of heavy metals and date rape drugs. PLOS ONE 14, e0210940 (2019).

[14] Watson, J. L. et al. De novo design of protein structure and function with rfdiffusion. Nature 620, 1089–1100 (2023).

[15] Amalfitano, E. et al. A glucose meter interface for point-of-care gene circuit-based diagnostics. Nat. Commun. 12, 724 (2021).

[16] Iyer, S., Karig, D. K., Norred, S. E., Simpson, M. L. & Doktycz, M. J. Multi-input regulation and logic with t7 promoters in cells and cell-free systems. PLoS ONE 8, e78442 (2013).

[17] Cress, B. F. et al. Rapid generation of crispr/dcas9-regulated, orthogonally repressible hybrid t7-lac promoters for modular, tuneable control of metabolic pathway fluxes inescherichia coli. Nucleic Acids Research 44, 4472–4485 (2016).

[18] Sakono, M. & Hayakawa, R. Repressor-like on-off regulation of protein expression by the dna-binding transcription activator-like effector in t7 promoter-based cell-free protein synthesis. ChemBioChem 22, 888–893 (2020).

[19] Kim, J.-S. & Pabo, C. O. Getting a handhold on dna: Design of poly-zinc finger proteins with femtomolar dissociation constants. Proceedings of the National Academy of Sciences 95, 2812–2817 (1998).

[20] Blackburn, M. C., Petrova, E., Correia, B. E. & Maerkl, S. J. Integrating gene synthesis and microfluidic protein analysis for rapid protein engineering. Nucleic Acids Res. 44, e68 (2016).

[21] Mandell, J. G. & Barbas, C. F., 3rd. Zinc finger tools: custom DNA-binding domains for transcription factors and nucleases. Nucleic Acids Res. 34, W516–23 (2006).

[22] Filonov, G. S., Moon, J. D., Svensen, N. & Jaffrey, S. R. Broccoli: Rapid selection of an rna mimic of green fluorescent protein by fluorescence-based selection and directed evolution. Journal of the American Chemical Society 136, 16299–16308 (2014).

[23] Hussey, B. J. & Moore, M. J. Programmable t7-based synthetic transcription factors. Nucleic Acids Research 46, 9842–9854 (2018).

[24] Ichikawa, D. M. et al. A universal deep-learning model for zinc finger design enables transcription factor reprogramming. Nature Biotechnology 41, 1117–1129 (2023).

[25] Marchand, A. et al. Targeting protein–ligand neosurfaces with a generalizable deep learning tool. Nature 639, 522–531 (2025).

[26] Li, Y. et al. Plasma concentrations of venetoclax and pharmacogenetics correlated with drug efficacy in treatment naive leukemia patients: a retrospective study. Pharmacogenomics J. 24, 37 (2024).

[27] Yasu, T., Gando, Y., Nomura, Y., Kosugi, N. & Kobayashi, M. Determination of venetoclax concentration in plasma using high-performance liquid chromatography. Journal of Chromatographic Science 61, 480–483 (2022).

[28] Mast, F. D. et al. Highly synergistic combinations of nanobodies that target sars-cov-2 and are resistant to escape. eLife 10 (2021).

[29] Gainza, P. et al. De novo design of protein interactions with learned surface fingerprints. Nature 617, 176–184 (2023).

[30] Pacesa, M. et al. Bindcraft: one-shot design of functional protein binders (2024). Preprint at 10.1101/2024.09.30.615802.

[31] McSweeney, M. A. et al. A modular cell-free protein biosensor platform using split t7 rna polymerase. Science Advances 11 (2025).

[32] Amalfitano, E. et al. A glucose meter interface for point-of-care gene circuit-based diagnostics. Nature Communications 12, 724 (2021).

[33] Grasemann, L., Lavickova, B., Elizondo-Cantú, M. C. & Maerkl, S. J. Onepot pure cellfree system. Journal of Visualized Experiments 1 (2021).

[34] Lavickova, B., Laohakunakorn, N. & Maerkl, S. J. A partially self-regenerating synthetic cell. Nature Communications 11 (2020).

