## Supplementary Information for "A T7 RNAP regulatory toolbox for cell-free network engineering and biosensing applications"

### 1 Supplementary Figures

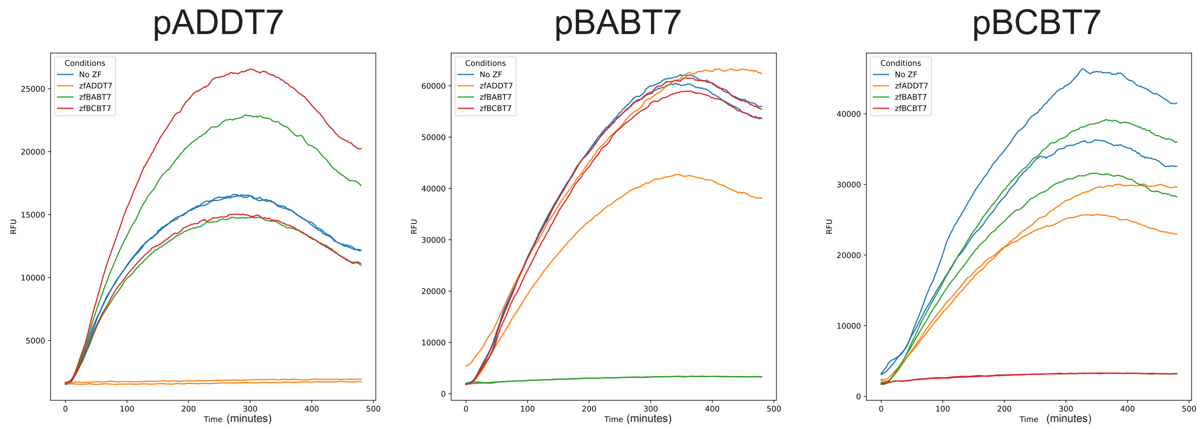


Figure S1: **Time-course of T7 promoter variants transcription under regulation of different repressors:** Time-course of transcription from the three different promoters in the presence of different ZF_PR-ZF_T7 repressors showing specific and potent repression by the matched repressor (n=2), the results show strong and specific repression with minimal cross-reactivity. The curves were processed by moving average with the window size of 5.

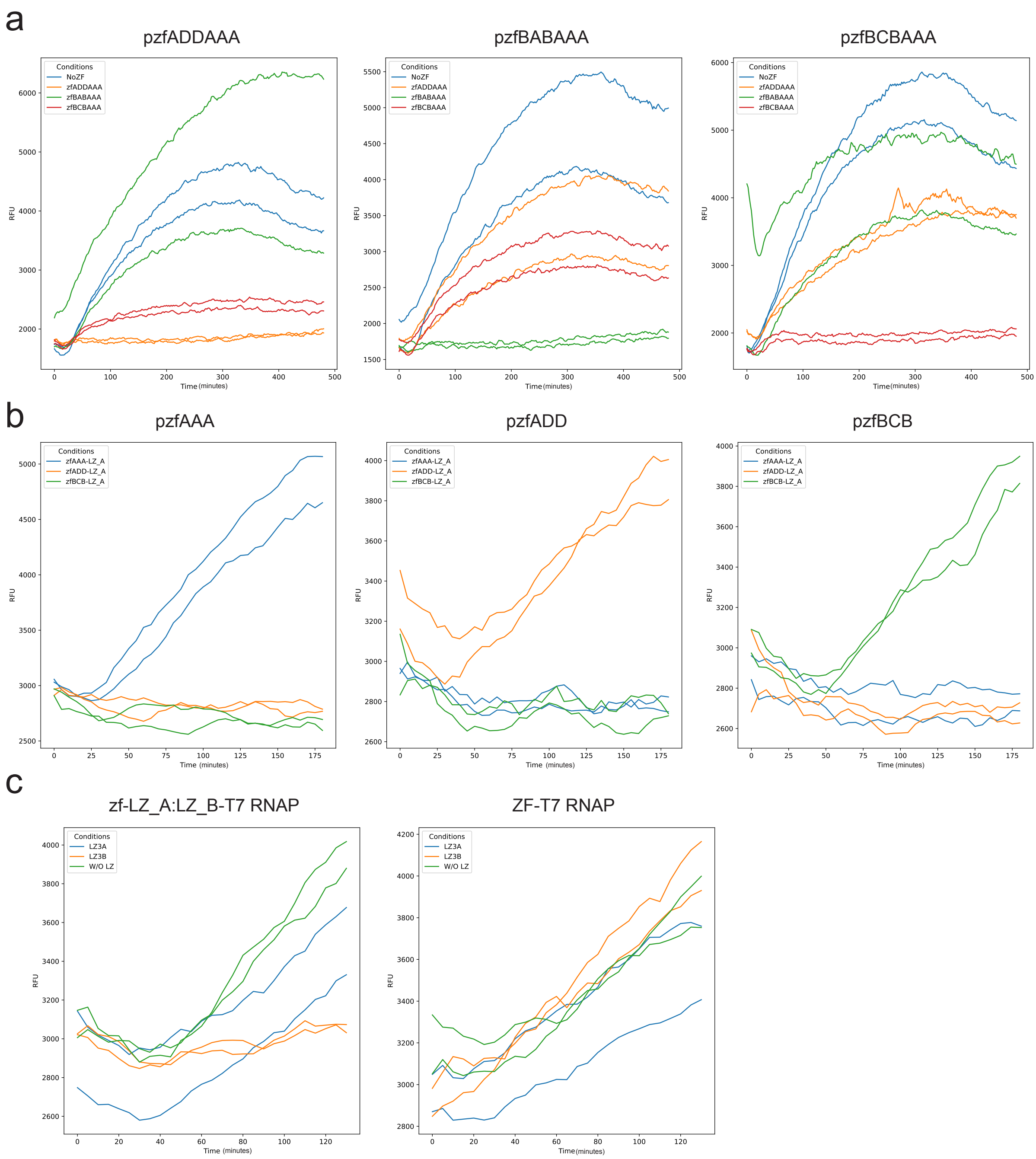


Figure S2: **Time-course transcription of regulatory component characterization:** **a,** Programmable pzfxxxAAAd1T7 promoter repression dynamic (n=2). The curves were processed by moving average with the window size of 5, which applies to all panels in this figure. **b,** Programmable pzfXXXd1T7 promoter activation dynamic (n=2). **c,** LZ repression (n=2), left panel is the result from LZ system that is repressible byLZ, the right panel if ZF-T7 RNAP system that won't be influenced by LZ.

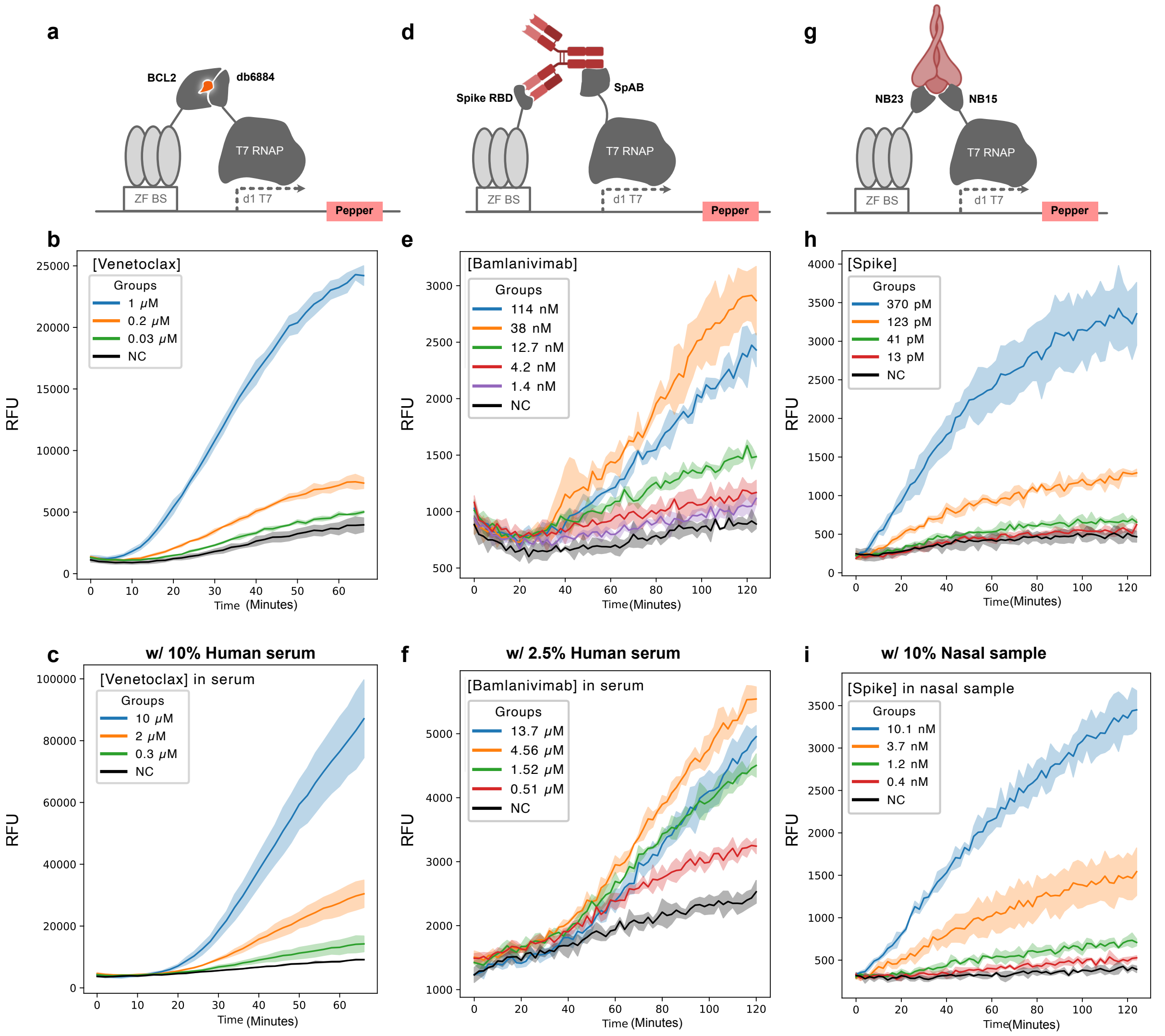


Figure S3: **Time-course of modular biomolecules sensor transcription:a,** Venetoclax biosensor. **b,** Venetoclax detection levels in a standard PURE reaction (n = 3; shaded regions indicate standard deviation; applies to all panels). **c,** Detection of Venetoclax in human serum. **d,** Antibody biosensor detecting Anti-Spike IgG. **e,** Detection of Bamlanivimab in a standard PURE reaction. **f,** Detection of Bamlanivimab in human serum. **g,** SARS-CoV-2 Spike protein biosensor. **h,** Detection of Spike in a standard PURE reaction. **i,** Detection of Spike in nasal swab sample.

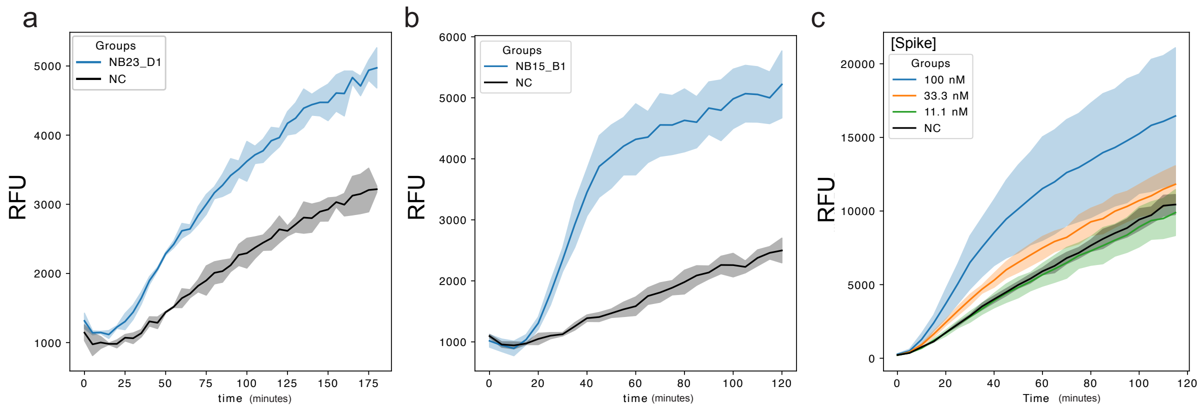


Figure S4: **Time-course of Spike detection reactions using de novo design binders:a,** Validation of ZF-NB23_D1 (n = 3; shaded regions indicate standard deviation; applies to all panels). **b,** Validation of ZF-NB15_B1. **c,** Spike detection with two de novo design binders complex NB23_D1 + NB15_B1- RNAP, demonstrating LOD of 33.3 nM.

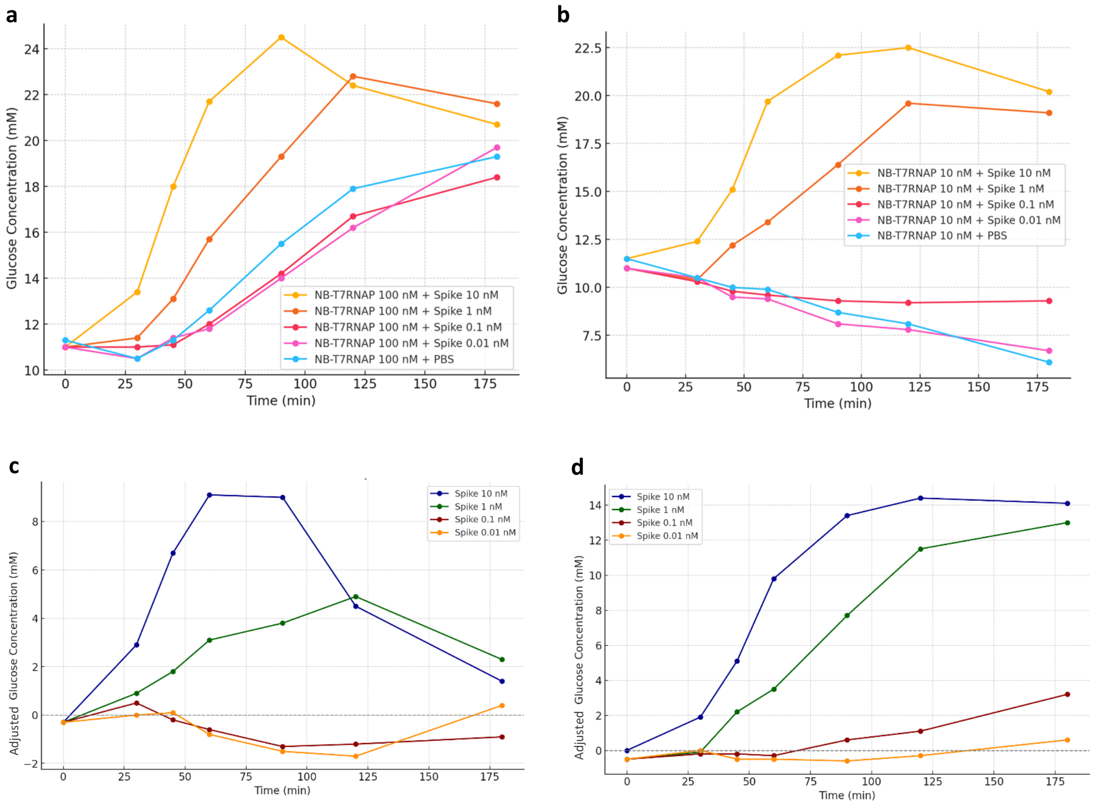


Figure S5: **Glucose signal over time for spike detection using △T7RNAP PURE with different NB15-T7RNAP concentrations:** Glucose concentration was measured at different time intervals in samples with varying spike concentrations diluted into PBS. **a**, Reaction with 100 nM NB15-T7RNAP **b,** Reaction with 10 nM NB15-T7RNAP. **c,** Result from panel a after adjusted glucose concentrations readout by subtracting the negative control signal. **d,** Result from panel b after adjusted glucose concentrations readout by subtracting the negative control signal. Each data point represents a single sample.

| | |
|---|---|
| Linear template structure | 5'common sequence - Promoter-5'UTR-RBS-ATG(start codon)-GOI- 3'UTR |
| 5'common sequence | GCTACTGATTGATGACGTGC |
| 5'UTR | ccacaacggtttccctctagaaataat |
| 3'UTR | cggctgctaacaaagcccgaaaggaagctgagttggc<br>tgctgccaccgctgagcaataactagcataaccccttggggcct<br>ctaaacgggtcttgaggggttttttgctgaaaggaggaactata<br>tcc |
| RBS | tttgtttaactttaagaaggagatatacc |

Table S1: **DNA template structure and sequences.** To design linear template for cell-free expression, follow the provided structure, then choose promoter and GOI sequence from following tables

| Promoter name | Sequence |
|---|---|
| WT T7 promoter | TAATACGACTCACTATAGGGAGA |
| pADDT7 | GCGGATGGAGTAATACGACTCACTATAGGGAGA |
| pBABT7 | GAGTGGGTGGTAATACGACTCACTATAGGGAGA |
| pBCBT7 | GAGGTAGTGGTAATACGACTCACTATAGGGAGA |
| pzf(ADDAAA)d1T7 | TCCATCCGCCGCCCACGCtgaattgatcGCTAGCCGACTCACTATAGGGAGA |
| pzf(BABAAA)d1T7 | CACCCACTCCGCCCACGCtgaattgatcGCTAGCCGACTCACTATAGGGAGA |
| pzf(BCBAAA)d1T7 | CACTACCTCCGCCCACGCtgaattgatcGCTAGCCGACTCACTATAGGGAGA |
| pzf(AAA)d1T7 | CGCCCACGCtgaattgatcGCTAGCCGACTCACTATAgggaga |
| pzf(ADD)d1T7 | TCCATCCGCtgaattgatcGCTAGCCGACTCACTATAGGGAGA |
| pzf(BCB)d1T7 | CACTACCTCtgaattgatcGCTAGCCGACTCACTATAgggaga |
| pzf(438)d1T7 | gtcctcactcggaattgatcGCTAGCCGACTCACTATAGGGAGA |
| zf AAA binding site | GCGTGGGCG |
| zf ADD binding site | GCGGATGGA |
| zf BAB binding site | GAGTGGGTG |
| zf BCB binding site | GAGGTAGTG |
| zf BCB binding site | GTAATACGA |

Table S2: **Promoter variants and ZF binding site sequence.**

| Name | Sequence |
|---|---|
| zfADDT7 | MHHHHHHGGSLEPGEKPYKCPECGKSFSQSGHLTEHQRTHTGEKPYKCPECGKSFS<br>QKSSLIAHQRTHTGEKPYKCPECGKSFSQSSSLVRHQRTHRQKDGGGSERPYACPV<br>ESCDRRFSDKTKLRVHIRIHTGQKPFQCRICMRNFSVRHNLTRHIRTHTGEKPFACD<br>ICGRKFARSDERKRHTKIHLRQ |
| zfBABT7 | MHHHHHHGGSLEPGEKPYKCPECGKSFSQSGHLTEHQRTHTGEKPYKCPECGKS<br>FSQKSSLIAHQRTHTGEKPYKCPECGKSFSQSSSLVRHQRTHRQKDGGGSERPY<br>ACPVESCDRRFSRNFILQRHIRIHTGQKPFQCRICMRNFSRSDHLTTHIRTHTG<br>EKPFACDICGRKFARHDQLTRHTKIHLRQ |
| zfBCBT7 | MHHHHHHGGSLEPGEKPYKCPECGKSFSQSGHLTEHQRTHTGEKPYKCPECGKS<br>FSQKSSLIAHQRTHTGEKPYKCPECGKSFSQSSSLVRHQRTHRQKDGGGSERPY<br>ACPVESCDRRFSRNFILQRHIRIHTGQKPFQCRICMRNFSQRSSLVRHIRTHTG<br>EKPFACDICGRKFARHDQLTRHTKIHLRQ |
| zfADDAAA | MHHHHHHGGSERPYACPVESCDRRFSDKTKLRVHIRIHTGQKPFQCRICMRNFSV<br>RHNLTRHIRTHTGEKPFACDICGRKFARSDERKRHTKIHLRQKDGGGSERPYAC<br>PVESCDRRFSRSDELTRHIRIHTGQKPFQCRICMRNFSRSDHLTTHIRTHTGEKP<br>FACDICGRKFARSDERKRHTKIHLRQ |
| zfBABAAA | MHHHHHHGGSERPYACPVESCDRRFSRNFILQRHIRIHTGQKPFQCRICMRNFSR<br>SDHLTTHIRTHTGEKPFACDICGRKFARHDQLTRHTKIHLRQKDGGGSERPYACP<br>VESCDRRFSRSDELTRHIRIHTGQKPFQCRICMRNFSRSDHLTTHIRTHTGEKPF<br>ACDICGRKFARSDERKRHTKIHLRQ |
| zfBCBAAA | MHHHHHHGGSERPYACPVESCDRRFSRNFILQRHIRIHTGQKPFQCRICMRNFSQ<br>RSSLVRHIRTHTGEKPFACDICGRKFARHDQLTRHTKIHLRQKDGGGSERPYACP<br>VESCDRRFSRSDELTRHIRIHTGQKPFQCRICMRNFSRSDHLTTHIRTHTGEKPF<br>ACDICGRKFARSDERKRHTKIHLRQ |
| zfAAA-LZ_A | MERPYACPVESCDRRFSRSDELTRHIRIHTGQKPFQCRICMRNFSRSDHLTTHIR<br>THTGEKPFACDICGRKFARSDERKRHTKIHLRQEIAALEQEIAALEKENAALEWE<br>IAALEQ |
| zfADD-LZ_A | MERPYACPVESCDRRFSDKTKLRVHIRIHTGQKPFQCRICMRNFSVRHNLTRHIR<br>THTGEKPFACDICGRKFARSDERKRHTKIHLRQEIAALEQEIAALEKENAALEWE<br>IAALEQ |
| zfBCB-LZ_A | MERPYACPVESCDRRFSRNFILQRHIRIHTGQKPFQCRICMRNFSQRSSLVRHIRT<br>HTGEKPFACDICGRKFARHDQLTRHTKIHLRQEIAALEQEIAALEKENAALEWEIAA<br>LEQ |
| eGFP | VSKGEELFTGVVPILVELDGDVNGHKFSVSGEGEGDATYGKLTLKFICTTGKLP<br>VPWPTLVTTLTYGVQCFSRYPDHMKQHDFFKSAMPEGYVQERTIFFKDDGNYKTR<br>AEVKFEGDTLVNRIELKGIDFKEDGNILGHKLEYNYNSHNVYIMADKQKNGIKVN<br>FKIRHNIEDGSVQLADHYQQNTPIGDGPVLLPDNHYLSTQSALSKDPNEKRDHM<br>VLLEFVTAAGITLGMDELYK |
| Trehalase | MGTAVRIDYASGLTDRENSMFKEIQLSGVFADSKTFVDSHPKLPLAEIAELYHVR<br>QQQAGFDLAAFVHRYFELPPSIASGFVSDTSRPVEKHIDILWDVLTRQPDRQEAG<br>TLLPLPYPYVVPGGRFREIYYWDSYFTMLGLQASKRWDLMEGMVNNFSHLIDTIG<br>FIPNGNRTYYEGRSQPPFYALMVELLANKQGESVLLAHLPHLRREYEFWMEGAAKL<br>SPAAPAHRRVVLLPDGSILNRYWDDIAAPRPESFREDYELAEAIGGNKRELYRHIR<br>AAAESGWDFSSRWFKDGNGMASIHTTDIIPVDLNALVFNLERMLAHIYGLQGDQDQ<br>ATHYYQLAEQRKQALLRYCWNAQQGFFHDYDYVAAQQTPVMSLAAVYPLYFSMVDQ<br>RTGDRVAEQIEAHFIQAGGVTTTLATTGQQWDAPNGWAPLQWLTIQGLRNYHHNSA<br>AEQIKQRWIALNQRVYRNTGKLVEKYNVYDLDVAGGGGEYELQDGFGWTNGVLLHL<br>LNESTP |
| NB15 | AQVQLVESGGGLVQAGGSLRLSCAASGRTFSSYAMGWFRQAPGKEREFVASINWNG<br>GNTYYADFVKGRFTISRDNAKNTVYLQMNSLKPEDTAVYYCAATGPNEYGLPREDL<br>FYDYWGQGTQVTVS |
| NB23 | AQVHLVESGGDLVQPGGSLRLSCVASGSGFENNAITWYRQAPGKERELVSGITSG<br>GSTNYADSVKGRFTISRDNAKNTVYLEMNSLKPEDTAVYLCQAVAWDSRRRSVVA<br>FWGQGTQVTVS |
| NB23_D1 | DKLKEKIKKLIQAIRDGASQDEIDKLLDEVAELGSGNLDLAILLNLITNLYLNG<br>YTFKEAEEAYKKVKAEKTTEEEKYLAEIYKNIKELLKKLGVDDDVF |

| Name | Sequence |
|---|---|
| NB15_C1 | MATIEIDIKNEKTGRQAYLRYNATPENADLVIDNAEADIAAFAKLDPGNTVSIR<br>GVADGGLEDLELARRAIERIEAAAKAAGVKVKSVELEASPSTQAAL |
| Spike_RBD | RVQPTESIVRFPNITNLCPFGEVFNATRFASVYAWNRKRISNCVADYSVLYNS<br>ASFSTFKCYGVSPTKLNDLCFTNVYADSFVIRGDEVRQIAPGQTGKIADYNYK<br>LPDDFTGCVIAWNSNNLDSKVGGNYNYLYRLFRKSNLKPFERDISTEIYQAGS<br>TPCNGVEGFNCYFPLQSYGFQPTNGVGYQPYRVVVLSFELLHAPATVCGPKKS<br>TNLVKNKCVNF |
| SpAB | ADNKFNKEQQNAFYEILHLPNLNEEQRNGFIQSLKDDPSQSANLLAEAKKLND<br>AQAPK |
| BCL2 | MAHAGRTGYDNREIVMKYIHYKLSQRGYEWDAGDDAEENRTEAPEGTESEVVHRA<br>LRDAGDDFERRYRRDFAEMSSQLHLTPDTARQRFETVVEELFRDGVNWGRIVAFF<br>EFGGVMCVESVNREMSPLVDNIAEWMTEYLNRHLHTWIQDNGGWDAFVELYGPSMR |
| DBVen1619 | MQYLLVVKGPNVTKFRWVDSSSEAETLARKIAKKLGLEVKSVEKKGNAVRVEI |
| ZF438-GSGGG-Binder | FQCRICMRNFSRQDRLDRHTRTHTGEKPFQCRICMRNFSQKEHLAGHLRTHTG<br>EKPFQCRICMRNFSRRDNLNRHLKTHGSGGG-Binder |
| Binder - T7 RNAP | MHHHHHH - Binder-<br>GSGGGGSGGGGSGGGGSMNTINIAKNDFSDIELAAIPFNTLADHYGERLAR<br>EQLALEHESYEMGEARFRKMFERQLKAGEVADNAAAKPLITTLLPKMIAR<br>INDWFEEVKAKRGKRPTAFQFLQEIKPEAVAYITIKTTLACLTSADNTTVQAV<br>ASAIGRAIEDEARFGRIRDLEAKHFKKNVEEQLNKRVGHVYKKAFMQVVEA<br>DMLSKGLLGGEAWSSWHKEDSIHVGVRCIEMLIESTGMVSLHRQNAGVVGQ<br>DSETIELAPEYAEAIATRAGALAGISPMFQPCVVPPKPWTGITGGGYWANGRR<br>PLALVRTHSKKALMRYEDVYMPEVYKAINIAQNTAWKINKKVLAVANVITKW<br>KHCPVEDIPAIEREELPMKPEDIDMNPEALTAWKRAAAAVYRKDKARKSRRI<br>SLEFMLEQANKFANHKAIWFPYNMDWRGRVYAVSMFNPQGNDMTKGLLT<br>LAKGKPIGKEGYYWLKIHGANCAGVDKVPFPERIKFIEENHENIMACAKSPLE<br>NTWWAEQDSPFCFLAFCFEYAGVQHHGLSYNCSLPLAFDGSCSGIQHFSAMLR<br>DEVGGRAVNLLPSETVQDIYGIVAKKVNEILQADAINGTDNEVVTVTDENTGEIS<br>EKVKLGTKALAGQWLAYGVTRSVTKRSVMTLAYGSKEFGFRQQVLEDTIQPAIDS<br>GKGLMFTQPNQAAGYMAKLIWESVSVTVVAAVEAMNWLKSAAKLLAAEVK<br>DKKTGEILRKRCAVHWVTPDGFPVWQEYKKPIQTRLNLMFLGQFRLQPTINTN<br>KDSEIDAHKQESGIAPNFVHSQDGSHLRKTVVWAHEKYGIESFALIHDSFGTIPA<br>DAANLFKAVRETMVDTYESCDVLADFYDQFADQLHESQLDKMPALPAKGNLNL<br>RDILESDFAFA |
| Broccoli_aptamer | ttgccatgtgtatgtgggagacggtcgggtccagatattcgtatctgtcgagtagagtgtgggctcccacat<br>actctgatgatccttcgggatcattcatggcaa |
| Pepper_aptamer | TTGCCATGTGTATGTGGGTTCGCCCACATACTCTGATGATCCCCAATCGTGGC<br>GTGTCGGCCTGCTTCGGCAGGCACTGGCGCCGGGATCATTCATGGCAA |

Table S3: **Sequence of proteins and reporters.**

| Components | Buffer A | Buffer B | Buffer HT | Stock buffer A | Stock buffer B |
|---|---|---|---|---|---|
| HEPES | 50 mM | 50 mM | 50 mM | 50 mM | 50 mM |
| Ammonium chloride | 1000 mM | - | - | - | - |
| Magnesium chloride | 10 mM | 10 mM | 10 mM | 10 mM | 10 mM |
| Potassium chloride | - | 100 mM | 100 mM | 100 mM | 100 mM |
| Imidazole (pH=7) | - | 500 mM | - | - | - |
| Glycerol | - | - | - | 30% (v/v) | 60% (v/v) |
| TCEP | 1 mM | 1 mM | 1 mM | 1 mM | 1 mM |

Table S4: **Protein purification Buffers.**

| Components | Concentration |
|---|---|
| HEPES | 250 mM |
| Potassium glutamate | 500 mM |
| Magnesium acetate | 59 mM |
| ATP | 5 mM |
| GTP | 5 mM |
| CTP | 5 mM |
| UTP | 5 mM |
| TCEP | 5 mM |
| Spermidine | 10 mM |

Table S5: **5x IVT buffer.**

| Components | volume or concentration |
|---|---|
| PURExpress Solution A | 4 µL |
| PURExpress Solution B | 3 µL |
| Spike protein | 10 nM |
| ZF-binder DNA template (0.1ng/µl) | 100 pg |
| NB15-T7RNAP or NB23-T7 RNAP | 10 nM |
| HBC 620 | 10 µM |
| Reporter DNA template | 10 nM |
| **Total Volume** | **10 uL** |

Table S6: **Composition of the cell-free reaction for de novo design binder screening.**

| Components | Without Amplifier | With Amplifier |
|---|---|---|
| PURExpress Solution A | 4 µL | 4 µL |
| Homemade ΔT7RNAP PURE | 1.3 µL | 1.3 µL |
| NEB Ribosome | 0.45 µL | 0.45 µL |
| ZF438-NB23 | 50 nM | 50 nM |
| NB15-T7RNAP | 10 nM | 10 nM |
| LZ_B-T7RNAP | – | 10 nM |
| NEB RNase Inhibitor | 10 U | 10 U |
| Trehalose | 10 mM | 10 mM |
| Template: pZFd1T7-ZF-LZ_A | – | 3 nM |
| Template: pZFd1T7-Tre37A | 20 nM | 20 nM |
| **Total Volume** | **10 uL** | **10 uL** |

Table S7: **Composition of the cell-free reaction for spike detection via glucose generation with and without amplifier.**

| Components | Without Amplifier | With Amplifier |
|---|---|---|
| PURExpress Solution A | 4 uL | 4 uL |
| Homemade ΔT7RNAP PURE | 1.3 uL | 1.3 uL |
| NEB Ribosome | 0.45 uL | 0.45 uL |
| ZF438-NB23 | 50 nM | 50 nM |
| NB15-T7RNAP | 10 nM | 10 nM |
| LZ_B-T7RNAP | - | 10 nM |
| NEB RNase Inhibitor | 10 U | 10 U |
| HBC620 | 10 nM | 10 nM |
| Template: pZFd1T7-ZF-LZ_A | - | 1 nM |
| Template: pZFd1T7-Pepper | 6 nM | 6 nM |
| **Total Volume** | **10 uL** | **10 uL** |

Table S8: **Composition of the cell-free reaction for spike detection using the Pepper aptamer, with and without amplifier.**
